# Cell-extrinsic controls over neocortical neuron fate and diversity

**DOI:** 10.1101/2024.10.15.618395

**Authors:** Natalia Baumann, Ilaria Morassut, Sergi Roig-Puiggros, Esther Klingler, Giorgia Bartolini, Sabine Fièvre, Denis Jabaudon

## Abstract

Cellular diversity in the neocortex emerges gradually during prenatal and postnatal development. While environmental interactions occur during this extended maturation period, the impact of extrinsic cues on determining the fate of distinct neuron types remains unknown. To address this question, we exposed developing neocortical cells to various environmental conditions and examined how this affects cell fate and diversity. Our developmental analyses reveal a hierarchical molecular program in which cell class-distinguishing features emerge first, followed by subclass- and type-related characteristics, with distinct developmental paces among cell populations. Environmental contribution was assessed in vivo, using genetically modified mice models in which position or innervation are altered, and in vitro using two-dimensional cultures. Acquisition of cellular identity and diversity remained stable across in vivo models. In contrast, in vitro glutamatergic neurons showed decreased expression of identity-defining genes, reduced diversity and alterations in canonical cortical connectivity. Cellular identity and diversity were restored towards in vivo values in organotypic slice cultures. These findings reveal cell population-specific responses to environmental conditions and highlight the role of extracellular context in shaping cell diversity in the maturing neocortex.

## Introduction

The neocortex consists of multiple cell populations whose identity can be defined by unique combinations of morphologies, spatial arrangements, connectivity, and gene expression (*1*). Understanding how neuronal identity emerges during development is important because transcriptional programs ultimately relate to specific connectivity and circuit function. The neocortex has a particularly prolonged developmental period compared to other brain structures: in many species it is still immature at birth, as evidenced by ongoing cell migration and myelinization, and incomplete circuit formation (*2*). Environmental conditions, including input-dependent factors, are thought to influence cell fate during critical periods of postnatal development (*3*), as some cortical neuron types appear particularly sensitive to external stimuli. For instance, layer (L) 2/3 neurons in the primary visual cortex need visual input to acquire their identity (*4*), L4 neurons in the primary somatosensory cortex require tactile stimulation to organize into whisker-related barrels (*5*), and some subtypes of GABAergic neurons depend on activity for migration (*6*). However, for most cell populations, the extent to which cell-intrinsic factors contribute to cell fate and diversity remains unknown. Such data is critical as many neuropsychiatric disorders, including autism spectrum disorder, are thought to emerge through complex interactions between genetic factors and the environment (*7–9*). A comprehensive evaluation of cell-extrinsic controls over cell-type-specific fate is thus essential and is the topic of the current study.

A major limitation in quantifying environmental effects across cell populations is the difficulty in defining and normalizing environment-induced responses across cell types. To address this limitation, here we first determined the transcriptional identities of mature cortical cells, which we then used to build a hierarchical reference framework to classify query cells following environmental manipulations. We used multiple paradigms to evaluate environmental impact on identity and diversity: in vivo studies in Reeler mouse mutants with abnormal cellular positioning (*10*) and VB^−^ mice with disrupted thalamocortical innervation (*5*), and in vitro investigations using two-dimensional cultures and organotypic slice cultures. We systematically examined how these distinct environmental conditions affected cell fate. While cellular identity and diversity remained stable in in vivo models, this stability was not maintained in vitro. Identity-defining transcriptional programs were particularly sensitive to in vitro conditions, leading to a loss of cellular diversity, with glutamatergic neurons being most affected. In organotypic slice cultures, cellular identity and diversity reached levels similar to those observed in vivo. Together, our findings uncover neocortical cell-population-specific sensitivities to extrinsic cues and underscore the critical role of the extracellular context in establishing cellular diversity during postnatal development.

## Results

### Cell population-specific maturation dynamics in the neocortex

Neocortical cells can be hierarchically categorized into *classes*, *subclasses*, and *types* (together, cell *populations*) based on their molecular identities (*1*). Classes are the broadest category, essentially distinguishing neuronal and non-neuronal cells, while types correspond to the finest-grained classification. To benchmark the developmental emergence of this organization, we first determined the mature hierarchy of cells in the adult mouse primary somatosensory cortex (SSp). Using unbiased iterative clustering of neocortical cells at incremental resolutions, we analyzed a comprehensive single-cell transcriptomic dataset (**Fig. 1A**, **Fig. S1A,B, Methods**) (*11*). We identified four cell classes, corresponding to glutamatergic neurons, GABAergic neurons, astrocytes and oligodendrocytes. At the next hierarchical level, five subclasses emerged, which, within glutamatergic neurons, distinguished intratelencephalic (IT) and extratelencephalic (ET) neurons. Finally, the highest resolution level corresponded to a repertoire of thirteen cell types, including eight types of glutamatergic neurons. At this clustering resolution, GABAergic neurons comprised two types, as did oligodendrocytes, while astrocytes belonged to a single population across hierarchical levels. The Oligo1 type expressed markers genes of oligodendrocytes precursor cells and maturing oligodendrocytes (e.g. *Pdgfrα*), while the Oligo2 type expressed markers of myelinating oligodendrocytes (e.g. *Mbp*) (**Table S1**). This does not imply that GABAergic interneurons, astrocytes, or oligodendrocytes are inherently less diverse than glutamatergic neurons (see e.g. reference *12*), but instead reflects the specific granularity of our analysis at the chosen time points. This iterative clustering technique showed similar grouping of cells when performed on our own P14 in vivo dataset, indicating a similar hierarchy of population at this stage of maturation (**Fig. S1B**).

**Figure 1.**
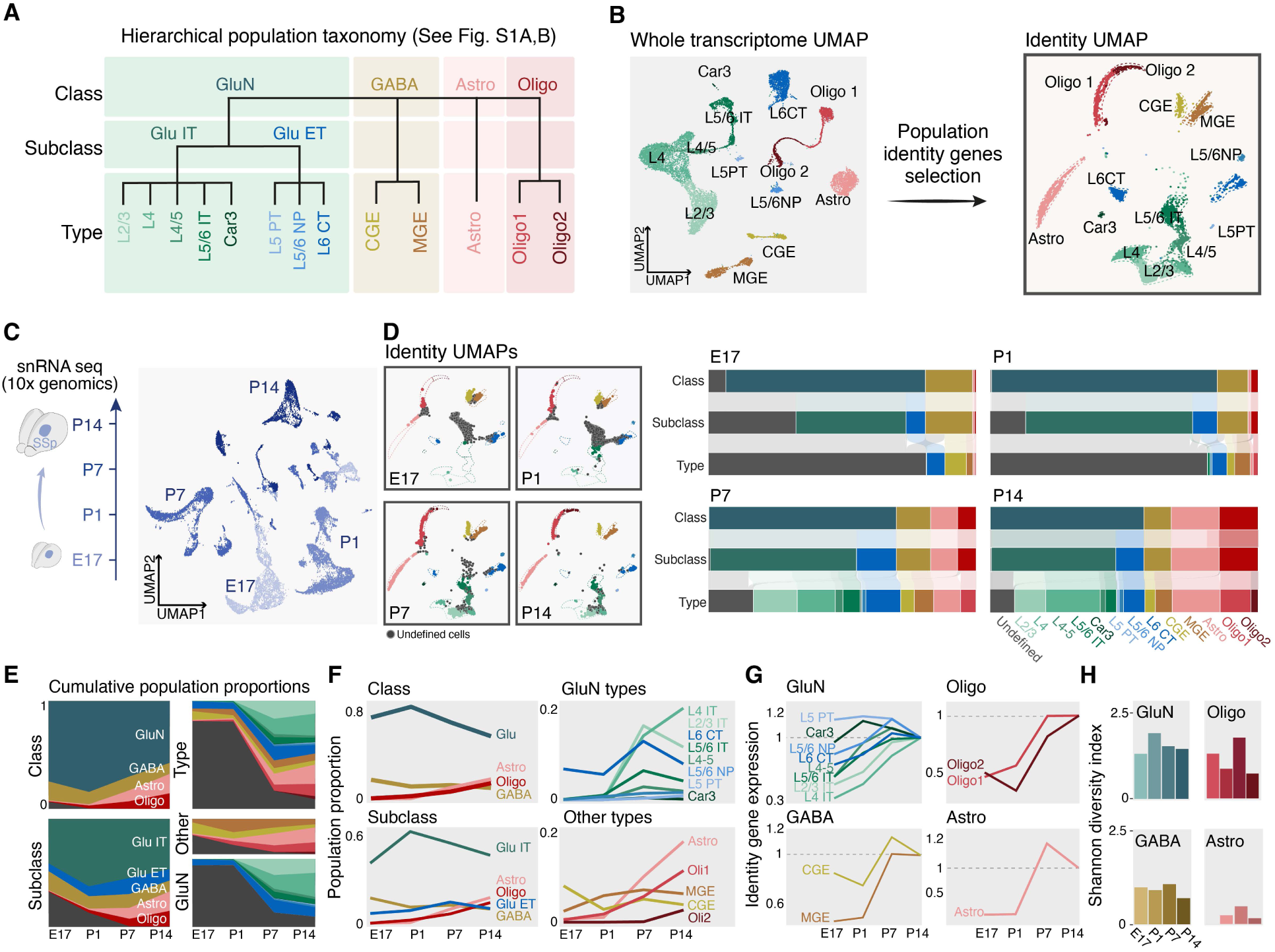
Cell population-specific maturation dynamics in the neocortex. **A,** Hierarchical tree containing four cell classes, five cell subclasses and thirteen cell types built by iterative clustering of the Allen Brain single-cell RNA sequencing dataset (Yao et al., 2021(*11*))(see. Fig. S1A,B). **B,** UMAP representation of the cell types of the reference dataset (left) and its associated identity UMAP calculated on population identity genes (right). The dotted lines around each cluster are used as a visual reference for comparison in the ID UMAP plots of all the following panels. **C,** Schematic of experimental design (left) and UMAP representing the sequenced nuclei at the indicated time points. **D,** Identity UMAPs at each indicated time point. Cell types color-coded as in A and B(left). Undefined cells, labeled grey, decrease as development progresses. The Sankey diagrams on the right summarize the relative proportions for each hierarchical level per time point and early emergence of class definition. **E,** Cumulative proportions (y axis) for classes and subclasses across the four time points analyzed (x axis) (left). On the right, the same type of plot is presented for types (top) and for glutamatergic neurons (bottom) separated from the other types (middle). **F,** Population proportions for classes and subclasses across the four time points analyzed (x axis) (left). On the right, the same type of plot is presented for glutamatergic types (top) and for other types (bottom) separately. **G,** Average relative expression of identity genes for corresponding cell classes across the four time points analyzed. **H,** the four bar plots display the values of the Shannon index for each cell class separately across the four time points analyzed. *Abbreviations*: SSp: primary somatosensory cortex; GluN, glutamatergic neurons; GABA, GABAergic neurons; Astro, astrocytes; Oligo, oligodendrocytes; IT, intratelencephalic neurons; ET, extratelencephalic neurons; L6 CT, layer 6 corticothalamic neurons; PT, pyramidal tract neurons; L2/3, layer 2/3; NP, near-projecting neurons; CGE, caudal ganglionic eminence; MGE, medial ganglionic eminence; ID, identity.

A cell’s transcriptome dynamically reflects its state, influenced by both intrinsic factors and extrinsic conditions. While some transcriptomic components are shared across populations, others are expressed specifically, constituting “identity”-defining transcriptional programs. To delineate these identity programs and distinguish features unique to specific cell populations, we identified differentially expressed genes at each hierarchical level (**Methods**). This allowed us to unbiasedly identify population “identity” genes, that categorize cells into classes, subclasses, and types (n = 1’566 class-specific transcripts, n = 1’534 subclass-specific transcripts, n = 3’452 type-specific transcripts, **Table S1**). Using these identity genes, we generated a corresponding “identity UMAP”, in which cells distribute according to their average expression of the identity genes (**Fig. 1B**, **Fig. S1C**). Within this reference space, query cells can be assigned a class, subclass, and type identity based on the identity of their nearest referenced neighbors or remain *undefined* when they do not reach a predefined threshold of proximity to single populations (**Fig. 1D**, **Fig. S1D, Methods**).

We assessed the developmental emergence of class/subclass/type hierarchy in this reference space. To this end, we microdissected the putative SSp on embryonic day (E)17.5, postnatal day (P) 1, P7 and P14, and performed single-nucleus RNA sequencing (snRNA-seq) (**Fig. 1C**). This strategy uncovered a hierarchical acquisition of mature cellular identities: classes were defined first, followed by subclasses and types. This maturation extended into the second postnatal week and was cell type-specific, with glutamatergic neurons reaching mature identities later than the other types (**Fig. 1D-F**). Glutamatergic neurons displayed type-specific developmental dynamics, with a sharp increase in IT neurons between P1 and P7, which was reflected in the expression dynamics of their identity-defining genes (**Fig. 1F,G**). Cellular diversity was highest in neurons compared to other classes, as measured using the Shannon index, which quantifies the evenness and richness of categories in a population (**Fig.1H**) (*13*). These data reveal a hierarchical molecular program governing cortical cell differentiation, characterized by distinct developmental paces among cell populations and their constituent types. In this program, class-distinguishing features emerge first, followed sequentially by subclass- and type-related characteristics.

### Cell type-specific environmental sensitivity in Reeler and VB^−^ mice

To examine environmental influences on cell fate, we used transgenic mouse models in which cellular environment is perturbed. To address the role of cellular position, we used Reeler mice, in which the laminar disposition of cells within the cortex is altered (*10*); to evaluate the role of synaptic input, we used VB^−^ mice, in which thalamic input to SSp is lacking (*5*)(**Fig. 2A**). We then analyzed the cell populations at each hierarchical level in these models and their respective controls using snRNA-seq (**Fig. 2B-D, Fig. S2A,B**). In both Reeler and VB-mice, all four classes, five subclasses, and thirteen types were present (**Fig. 2E; Fig. S2C-D**). Both mutants displayed more cells with *undefined* identities, predominantly positioned between L2/3 IT and L4 IT neuron clusters (**Fig. 2F**). This was more pronounced in VB^−^ mice and suggests a particular sensitivity of L2/3 IT and L4 IT neurons to input-dependent cues. VB^−^ mice displayed a shift in the proportion of these neuron types, with an increase in L2/3 IT neurons at the expense of L4 IT neurons (**Fig. 2G**). This finding, coupled with the reduced definition of L4 IT identity in both mutants (**Fig. 2H**), aligns with previous research suggesting an input-dependent emergence of L4 IT neurons from L2/3 IT-like neurons in SSp (*14*).

**Figure 2:**
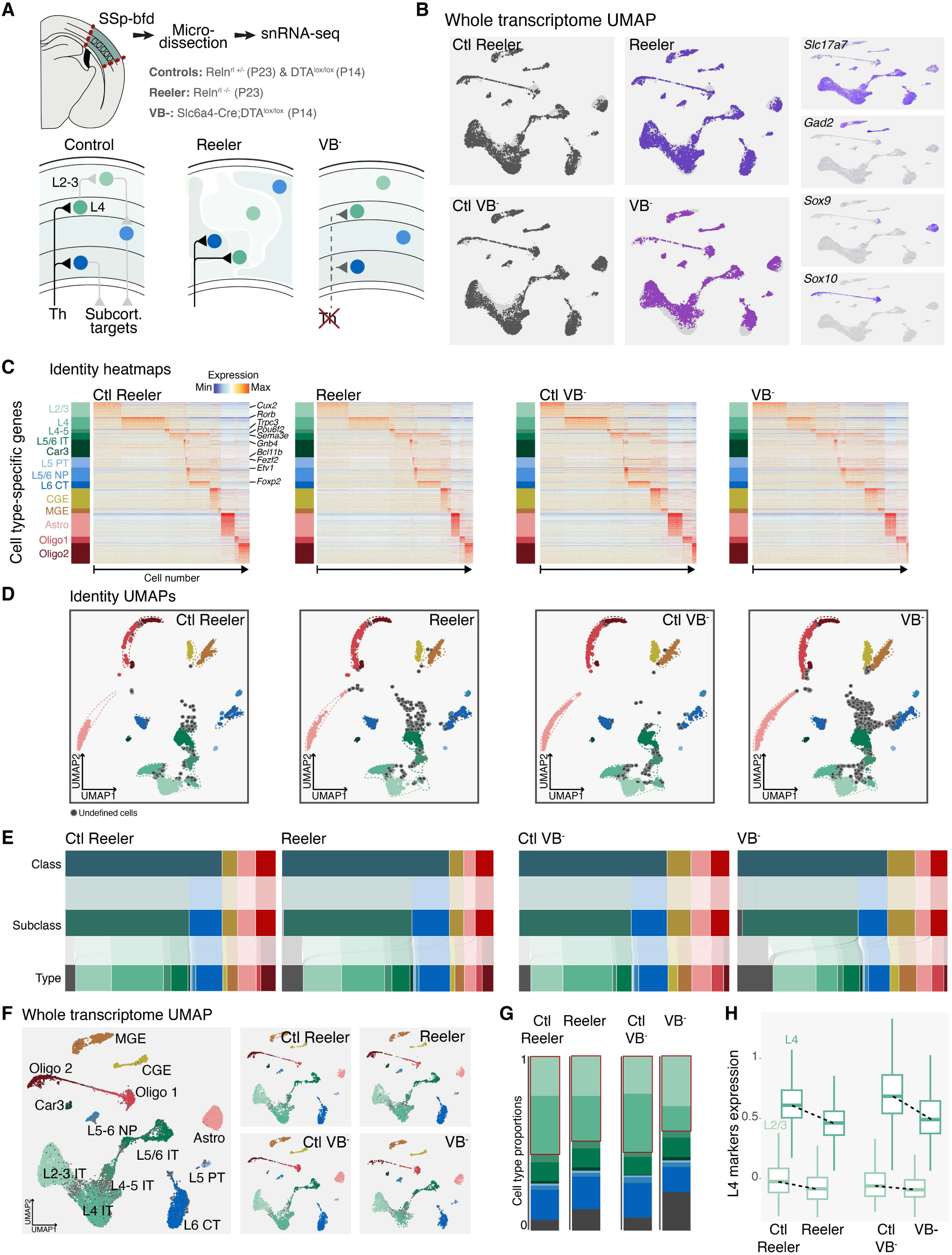
Cell type-specific environmental sensitivity in Reeler and VB^-^ mice. **A,** Schematic representation of snRNA-seq dataset generation for P23 Reeler and P14 VB^-^ mice model, and their respective controls. **B,** UMAP representation of snRNA sequencing datasets color-coded per condition using whole transcriptome data. On the right, feature plots for four class-specific markers are displayed. **C,** Heatmaps representing the cell type identity gene expression in Reeler and VB^-^ mice, and their respective controls. **D,** Identity UMAPs representing cell type classification for Reeler model and its respective control (left), and for VB^-^ and its respective control (right). Undefined cell types are color-coded in dark grey. The dotted lines around each cluster are used as a visual reference for comparison in all ID UMAP plots. **E,** Sankey diagrams showing the relative proportions of the cellular populations identified at each of the three hierarchical levels, represented separately for each of the analyzed conditions. **F,** Whole transcriptome UMAP for all conditions together (left) and split per condition (right). **G,** Proportions of glutamatergic neuron types across the four analyzed conditions. The red rectangles highlight the proportion of L2/3 and L4 IT types. **H,** Boxplots showing expression levels of L4 marker genes within L2/3 IT neurons (light green) and L4 IT neurons (dark green) across the four conditions. *Abbreviations*: SSp-bfd: primary somatosensory cortex, barrel fields; Th, thalamus; Ctl, control; Astro, astrocytes; Oligo, oligodendrocytes; IT, intratelencephalic neurons; L6 CT, layer 6 corticothalamic neurons; PT, pyramidal tract neurons; L2/3, layer 2/3; NP, near-projecting neurons; CGE, caudal ganglionic eminence-derived interneurons; MGE, medial ganglionic eminence-derived interneurons.

Interestingly, while these specific neuron types showed marked sensitivity to environmental changes, overall glutamatergic neuron diversity remained stable (**Fig. S2E**).

### Cell population-specific molecular responses in two-dimensional cultures

In vivo, compensatory mechanisms may mask potential environmental effects on cellular identity. To uncover such effects, we turned to an in vitro model. We assessed emergence of cellular identities with snRNA-Seq in 2D cultures of E16 dissociated SSp at day in vitro (DIV) 1, 4, 10 and 17, comparing to corresponding ages in vivo, i.e. E17 (corresponding to E16 + 1 day), P1 (E16 + 4 days), P7 (E16 + 10 days) and P14 (E16 + 17 days) (**Fig. 3A-C**). In this environment, all cell classes were able to emerge (**Fig. 3B**), with neurons exhibiting normal resting membrane potential and action potential firing, suggesting adequate culture conditions (**Fig S3A**).

**Figure 3:**
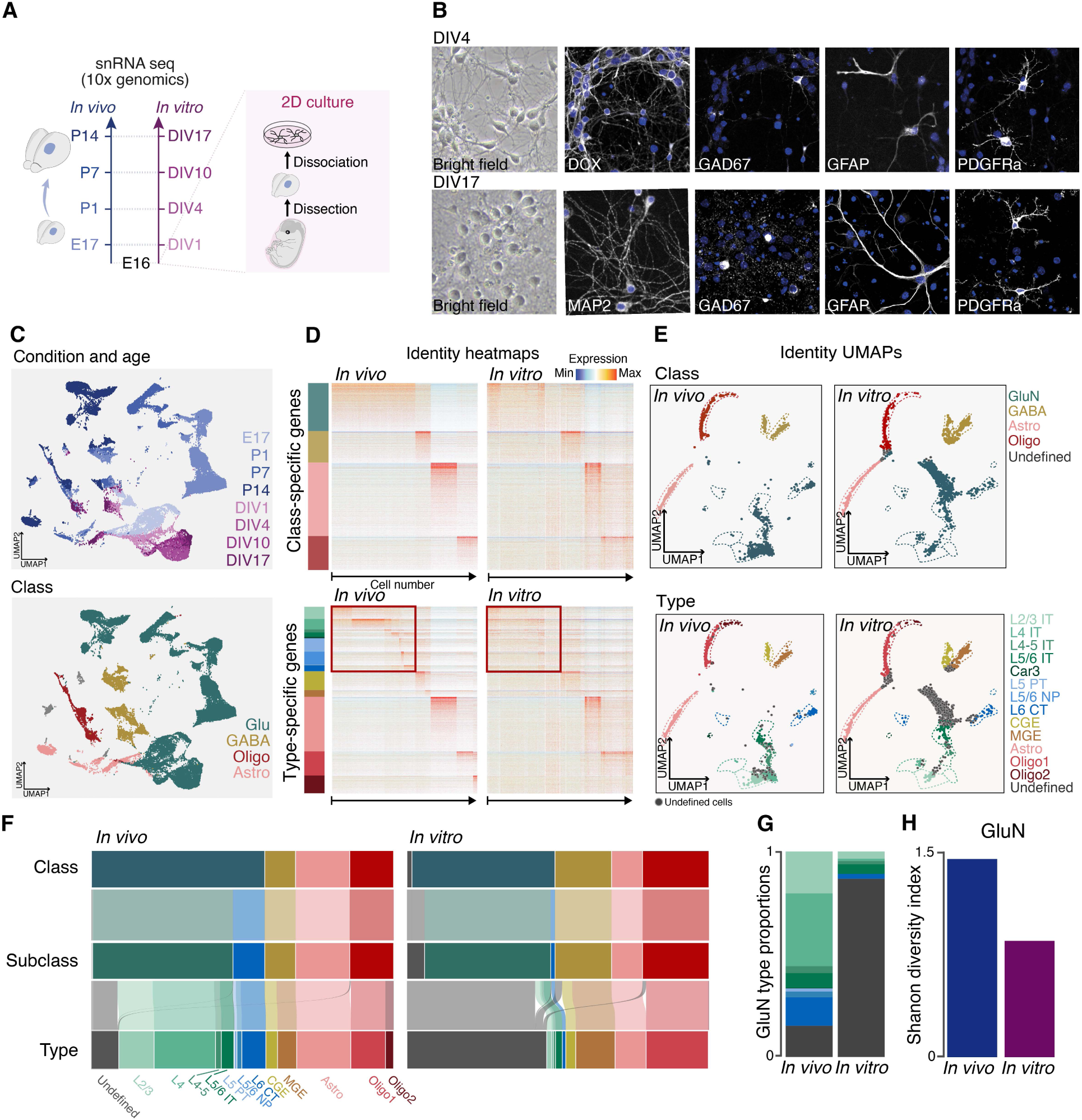
Cell population-specific molecular responses in 2D cultures. **A,** Experimental design of single nucleus sequencing of *in vitro* and *in vivo* samples. **B,** Left, representative bright field images of DIV4 and DIV17 cultures. Right, immunofluorescence staining of DIV4 and DIV17 cultures with glutamatergic (DCX, MAP2), GABAergic (GAD67), astrocytic (GFAP) and oligodendrocyte precursor (PDGFRα) markers. Scale bar: 50 µm. **C,** UMAP representation of snRNA sequencing dataset color-coded by condition timepoint (top) and by cell class annotation (bottom). **D,** Heatmaps displaying expression of identity makers for cell class and type genes (rows) for each cell (columns), *in vivo* (left) and *in vitro* (right). The red boxes indicate the genes that are most differently expressed between the two conditions. **E,** Identity UMAPs color-coded for cell classes (top) and types (bottom), for *in vivo* condition (left) and *in vitro* one (right). The dotted lines around each cluster are used as a visual reference for comparison in all ID UMAP plots. **F,** Sankey diagrams showing the relative proportions of cell populations at each hierarchical level, between *in vivo* at P14 (left) and *in vitro* at DIV17 (right). **G,** Relative proportions of glutamatergic neuron types, including undefined types (in grey), between *in vivo* at P14 and *in vitro* at DIV17. **H,** Shannon index for *in vivo* and *in vitro* glutamatergic neurons. *Abbreviations*: DIV, days in vitro; GluN, glutamatergic neurons; GABA, GABAergic neurons; Astro, astrocytes; Oligo, oligodendrocytes; Glu IT, glutamatergic intratelencephalic neurons; L6 CT, layer 6 corticothalamic neurons; PT, pyramidal tract neurons; L2/3, layer 2/3; NP, near-projecting neurons; CGE, caudal ganglionic eminence-derived interneurons; MGE, medial ganglionic eminence-derived interneurons; ID, identity.

In vitro class identities were similar to those observed in vivo (**Fig. 3C,D**). Relative proportions of classes were, however, altered, with a decrease in glutamatergic neuron fraction (**Fig. 3E,F**) suggesting that this class is more dependent on environmental cues for their survival or differentiation. All five subclasses were well delineated (**Fig. S3B,C, Fig. 3F**), but the proportion of ET glutamatergic neurons was strongly reduced. To assess ET neuron survival, we labeled ET and IT neurons at their birth dates, using in utero electroporation of GFP and TdTomato at E12.5 and E14.5, respectively. Comparable survival rates in vitro suggested that reduced ET neurons values reflect altered fate rather than selective death (**Fig. S3D, Methods**).

At the type level, however, many cells were classified as *undefined*, primarily within the glutamatergic class (**Fig. 3E-G**). This reflected a lower molecular definition of cellular identity at the type level, rather than changes in shared transcriptional programs (**Fig. 3D**). Glutamatergic types showed a lower definition of cellular identity, suggesting that *undefined* identities result from incomplete acquisition rather than a mixing of identities. Accordingly, expression of type markers was specifically decreased in glutamatergic neurons, while the gene expression of other types appeared less affected (**Fig. S3E)**. Consequently, glutamatergic neuron diversity was decreased (**Fig. 3H).** Non-neuronal cells, on the other hand, maintained their identity definition and diversity (**Fig. S3F**). Whole transcriptome analysis of differentially expressed genes between in vivo and in vitro classes revealed mostly class-specific changes (**Fig. S3G** left). In glutamatergic neurons, most environment-dependent genes coded for proteins associated with synaptic transmission (132 genes), cell-cell communication (81 genes) and ion transport (165 genes) (**Fig. S3G** right, **Table S2**). Hence, cell-extrinsic responses are class-specific.

To assess maturation pace in vitro, we applied an ordinal regression model trained on temporally dynamic in vivo genes. All classes showed similar maturation delays in vitro (**Fig. S3H, Table S3**. This suggests that observed differences in identity definition between classes do not primarily reflect differences in cellular maturity. Similarly, *undefined* cells within the glutamatergic neuron class had a comparable maturation stage than their defined counterparts (**Fig. S3I**). Together, these results reveal cell type-specific extrinsic controls over glutamatergic neuron identity and diversity that act largely independently of differentiation programs at these developmental stages.

### Canonical circuit connectivity is affected in 2D cultures

To understand how these molecular changes might affect neural circuit formation, we next examined the impact on specific neuron types and their connectivity patterns. IT neurons comprise distinct types that assemble to form intracortical circuits: L4 IT neurons receive thalamic input and forward it locally to L2/3 IT neurons, which in turn project to other L2/3 IT neurons. L4 IT to L2/3 IT neuron connectivity is unidirectional: while L4 IT neurons form synapses with L2/3 IT neurons, the reverse is rare. This arrangement ensures the forward flow of sensory information (**Fig. 4A**, left) (*15*).

**Figure 4:**
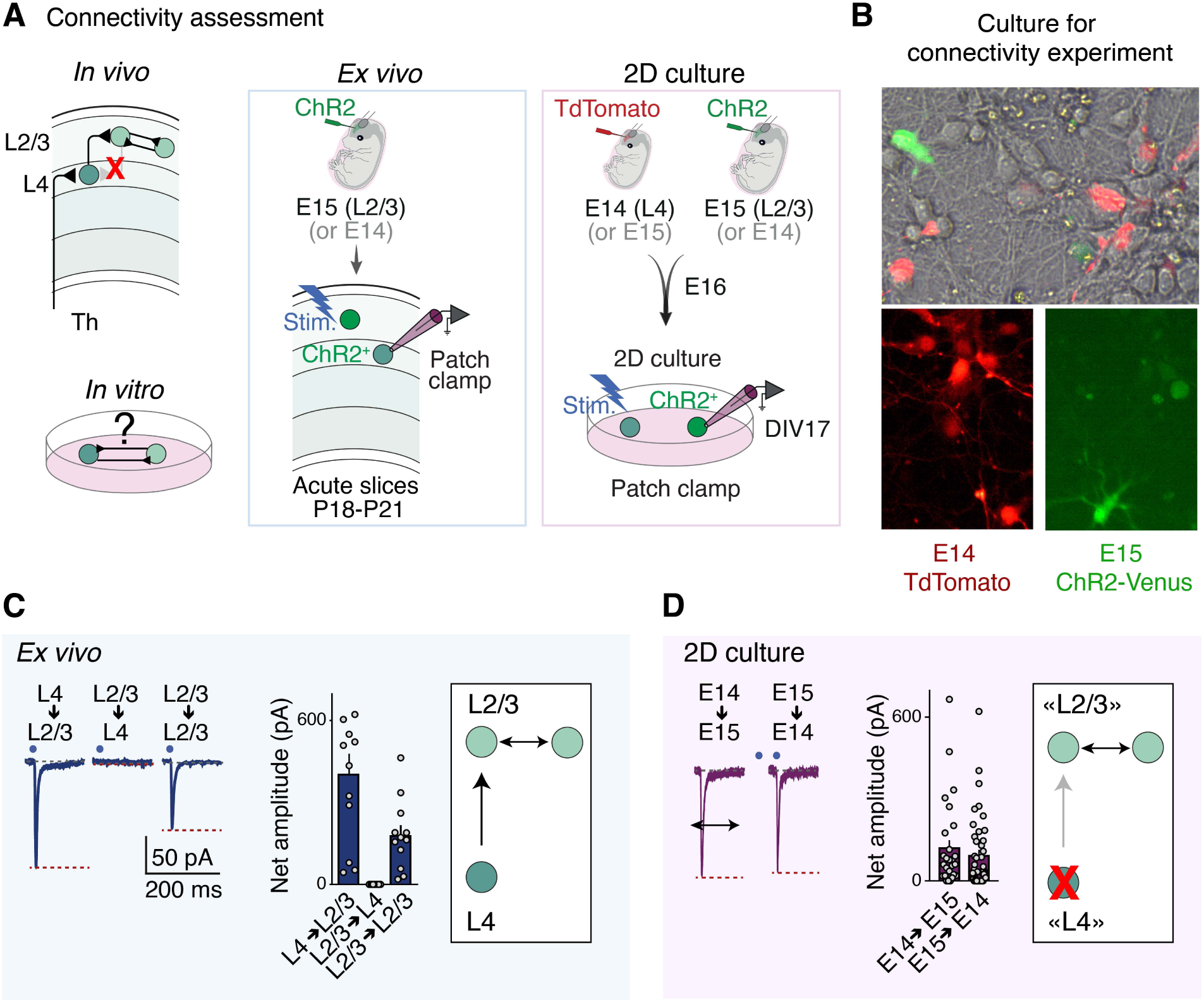
Canonical circuit connectivity is affected in 2 dimensional cultures. **A,** Schematic of the experiment used for connectivity assessment in vivo and in vitro. **B,** Example of immunofluorescence detected in the mixed cultures used for the connectivity experiment. **C,** Example of current traces obtained for each type of connectivity assessment (left), bar plot of net amplitudes measured for each condition (middle), and schematic of the circuit connectivity measured in vivo at P18-P21. **D,** Example of current traces obtained for each type of connectivity assessment (left), bar plot of net amplitudes measured for each condition (middle), and schematic of the circuit connectivity measured in vitro at DIV17 (right). *Abbreviations*: DIV, days in vitro; Stim., light stimulation; ChR2, channel rhodopsin 2; pA, pico amperes; ms, milliseconds.

L4 IT neurons were not found in vitro, suggesting that the identity of this cell type is highly dependent on extrinsic cues (**Fig. 2E**, **Fig. 3F-G**). To assess whether the lack of L4 IT neuron identity entailed a corresponding loss in unidirectional L4 IT-to-L2/3 IT connectivity, we targeted these two populations by in utero electroporation at E14 (to target prospective L4 IT neurons) and E15.5 (to target prospective L2/3 IT neurons) (**Fig. 4A**, right). To assess L4 IT-to-L2/3 IT connections, L4 IT neurons were targeted with channel rhodopsin (ChR2) at E14, while postsynaptic L2/3 IT neurons were targeted with a reporter gene (TdTomato) at E15.5 (see **Methods, Fig. 4B**). Conversely, to assess L2/3 IT-to-L4 IT connections, ChR2 was expressed in L2/3 IT neurons and TdTomato in L4 IT neurons. We shone blue light to stimulate ChR2-expressing neurons while recording evoked responses in postsynaptic patch-clamped Td-Tomato-expressing neurons. Ex vivo, light-stimulation of L4 IT neurons triggered a response in postsynaptic L2/3 IT neurons; L2/3 IT neurons did not evoke responses in L4 IT neurons, consistent with a unidirectional L4-to-L2/3 connectivity (**Fig. 4C**). In vitro, however, a reciprocal connectivity was found between putative L4 IT neurons and L2/3 IT neurons (**Fig. 4D**), as normally found between L2/3 IT neurons in vivo (**Fig. 4C**). Hence, environment-dependent changes in cell fate are not merely isolated transcriptional events, but instead have functional implications for the circuit properties of affected neurons.

### Increased cell type-specific glutamatergic neuron identity definition in organotypic cultures

To further explore the influence of cell-extrinsic cues on cell fate and diversity, we transitioned to organotypic slice cultures. These slices offer a more complex extracellular environment compared to 2D cultures, while maintaining the accessibility of an in vitro system. We hypothesized that this enhanced environment could restore some of the defects observed above. To test this, we cultured 300 µm-thick coronal slices of E16 embryonic somatosensory cortex, using the same medium as before (**Fig. 5A, B, Methods**). We performed snRNA-Seq at DIV10 and DIV17 (**Fig. 5C**) and analyzed identity and diversity at each level of the cellular hierarchy. At the class and subclass levels, the distribution of cellular populations in organotypic cultures more closely resembled in vivo distributions, contrasting with observations in 2D cultures. This included the restoration of ET neuron proportions. A larger fraction of glutamatergic neurons acquired type-specific identities, though not fully matching in vivo levels (**Fig. 5D-F, S4A,B**). Correspondingly, glutamatergic neuron type diversity approached in vivo values (**Fig. 5G**). The molecular definition of glutamatergic neuron identity was higher than in 2D cultures (**Fig. 5H** left). This was evident in the expression of key transcripts: IT neuron markers *Cux2* and *Rorb* (*16, 17*) and ET neuron markers *Bcl11b* and *Foxp2* (*18, 19*) were more highly expressed in organotypic cultures compared to 2D cultures (**Fig. 5H** right). More broadly, the average expression level of cell type-specific markers increased to levels observed in vivo for almost all cell populations (**Fig. S4C**).

**Figure 5:**
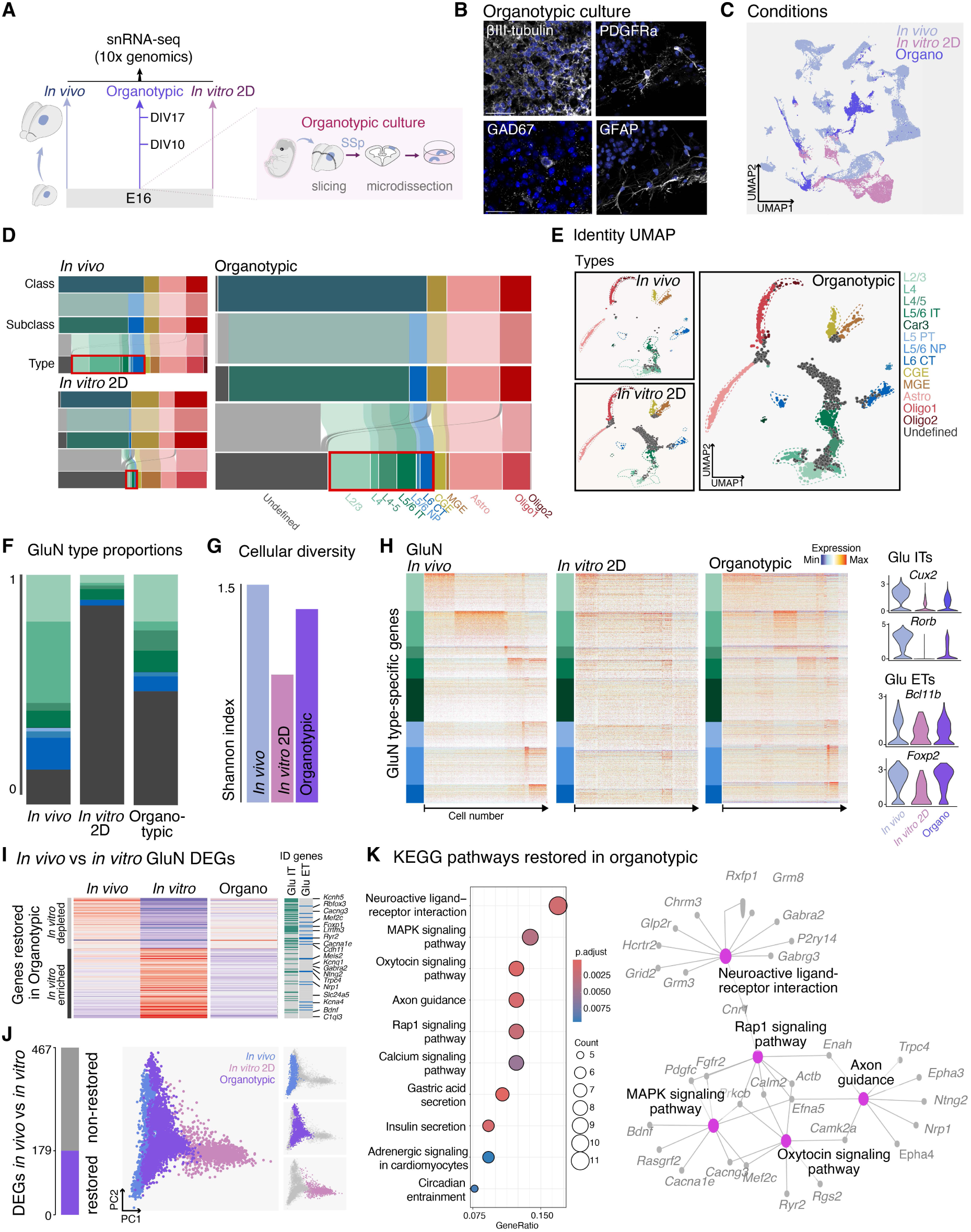
Increased cell type-specific glutamatergic neuron identity definition in organotypic slices. **A,** Schematic of the experimental pipeline, and of organotypic culture method (right). *In vivo* and 2D primary culture data are the same that were used previously. **B,** Immunofluorescence images of DIV17 organotypic cultures for representative markers of glutamatergic neurons (βIII-tubulin), GABAergic neurons (GAD67), oligodendrocyte precursors (PDGFRα), astrocytes (GFAP). Scalebar: 50µm. **C,** UMAP of snRNA-seq dataset now including *in vivo*, organotypic culture and *in vitro* 2D primary culture cells. **D,** Sankey diagrams for all populations and cumulative proportions for glutamatergic types only, for all three conditions, are displayed as in Fig. 3, with the new data for organotypic culture-derived cells. **E,** Identity UMAPs for cell types are displayed for *in vivo* and *in vitro* 2D (same data as in Fig. 3), and for organotypic culture (new data). Undefined types are color-coded in grey. The dotted lines around each cluster are used as a visual reference for comparison in all ID UMAP plots. **F,** Proportions of glutamatergic neuron types are displayed for *in vivo* and *in vitro* 2D (same data as in Fig. 3), and for organotypic culture (new data). Undefined types are color-coded in grey. **G,** Shannon index for cellular diversity within glutamatergic neurons across the three conditions. **H,** Identity heatmaps only for glutamatergic neuron genes are displayed for *in vivo* and *in vitro* 2D (same data as in Fig. 3), and for organotypic culture (new data). Expression levels for selected genes are shown on the right. **I,** Top, heatmap showing the average expression level of differentially expressed genes (rows) in vivo compared to in vitro, which were restored in organotypic culture in glutamatergic neurons. The values are displayed for each condition at the latest timepoint (P14 in vivo and DIV17 in in vitro 2D and organotypic). ID genes of IT and ET glutamatergic neurons are reported on the right. **J,** PCA analysis of average expression of the restored genes in organotypic culture for each cell (dots), color-coded per condition. On the right, individual panels show single conditions. **K,** KEGG enrichment pathway analysis of genes whose expression was rescued in glutamatergic neurons cultured in organotypic slices (left). Cnet plot indicating the genes involved in the top 5 pathways is reported (right). *Abbreviations*: GluN, glutamatergic neurons; IT, intratelencephalic neurons; ET, extratelencephalic neurons; Astro, astrocytes; Oligo, oligodendrocytes; Organo, organotypic; L6 CT, layer 6 corticothalamic neurons; PT, pyramidal tract neurons; L2/3, layer 2/3; NP, near-projecting neurons; CGE, caudal ganglionic eminence-derived interneurons; MGE, medial ganglionic eminence-derived interneurons; ID, identity.

To assess whether the increased molecular definition of cellular identity in organotypic conditions reflected effects on cellular maturation, we performed pseudo time analysis (**Fig. S4D**). This analysis revealed comparable maturity levels in both 2D cultures and organotypic cultures. This suggests that the enhanced molecular definition of cellular identity observed in organotypic cultures is largely independent of overall maturation levels.

To identify the transcriptional programs at play in restoring glutamatergic neuron type-specific identities, we identified differentially expressed genes between in vivo and 2D-cultured neurons at P14 and DIV17 (467 genes). This analysis revealed a subset of genes (179) with similar patterns of expression in vivo and in organotypic slices, compared to 2D cultures, i.e. “in vivo-like” genes (**Fig. 5I, Fig. S4E**). Of all the genes that were different, 38% were “rescued”. Their average expression in organotypic cultures closely recapitulates the expression pattern of in vivo cells, as revealed by PCA analysis (**Fig. 5J, Fig S4F**). These genes included IT and ET type-specific markers (e.g., *Rbfox3*, *Foxp1*; **Fig. 5I** right) and genes involved in extrinsic regulation, such as those associated with neuroactive ligand-receptor interaction, axon guidance, and MAPK signaling pathways (e.g., *Epha3*, *Nrp1*; **Fig. 5K**), as identified by KEGG pathway analysis. Notably, the MAPK signaling pathway plays crucial roles in cortical development through extrinsic modulation, and its dysregulation has been linked to neurodevelopmental conditions such as autism spectrum disorder(*20, 21*). These findings suggest that organotypic cultures preserve key extrinsic cues necessary for establishing and maintaining cell type-specific molecular identities.

## Discussion

Our comprehensive analysis of neocortical cellular identity across various environmental conditions reveals cell type-specific extrinsic influences on neuronal fate. Using a bioinformatic framework based on population-specific cellular identity, we report a remarkable robustness of class and subclass identity, which remained conserved even in 2D dissociated cultures. In vivo, acquisition of cell type identity was only minimally affected by changes in cellular position (Reeler mice) or input (VB^−^ mice). In Reeler mice, however, the distribution of cells across canonical types was affected, as evidenced by an increase in L2/3 IT neurons at the expense of L4 IT neurons. This could reflect an incomplete maturation of L4 IT neurons, which initially send long-range axons to the opposite hemisphere (*14*). Since L4 IT identity is not fully defined before P14, these neurons may thus initially have a L2/3 IT-type molecular identity that normally matures to become L4 IT in an input-dependent manner.

Our in vitro experiments reveal a neuron class-specific susceptibility to extrinsic cues for identity acquisition, with glutamatergic neurons being more susceptible than GABAergic neurons. This difference may reflect the resolution thresholds used to define cell types in this study, as some subtypes of GABAergic neurons (not studied specifically here) have been shown to mature differently in different environments (*22, 23*). Similarly, astrocytes change their properties in vitro (*24*). This is reflected in our 2D cultures by an increase in the number of differentially expressed genes in vitro. Finally, oligodendrocyte maturation highly depends on culture conditions (*25, 26*), which is reflected here by a specific decrease of the mature oligodendrocytes type (Oligo2).

The improved molecular definition of cellular identity in organotypic slice cultures underscores the crucial role of physiological environments in determining neuronal fate. This finding supports the significance of extrinsic modulation in cell differentiation and highlights the potential of advanced 3D culture systems, such as organoids, in replicating complex neural tissues. Our results suggest that manipulating the cellular microenvironment could be used to direct cell fate in stem cell-based therapies. Further investigation into these environment-dependent mechanisms may identify novel therapeutic targets in neurodevelopmental disorders, particularly those influenced by gene-environment interactions.

## Methods

### Experimental Models

All experimental procedures were approved by the Geneva Cantonal Veterinary Authority. Embryonic day (E) 0.5 was established as the day of vaginal plug. Wild-type CD1 mice were provided by Charles River Laboratories.

Male and female embryos between E12.5 and E15.5 were used for the in utero electroporation experiments, and mice between E16 and P21 for the postnatal experiments. Pregnant dams were kept in single cages and pups were kept with their mothers until P21, in the institutional animal facility under standard 12:12h light/dark cycles.

Reeler mice (https://www.jax.org/strain/000235) were obtained from Prof. Jochen F. Staiger (University Medical Center Göttingen, Centre for Anatomy, Institute of Neuroanatomy, Kreuzbergring 36, D-37075, Göttingen). Mutant mice were obtained by crossing Reln^rl^/+ heterozygous mice. The line was genotyped using the following primers: CTGTCCTCACTCTGCCCTCTGGTAG; CTACACAGTTGACATACCTTAATCTAC; ACTTGCATTAATGTGCAGTGTTGTC.

Vb- mouse model was obtained by crossing Slc6a4-Cre transgenic mice (*27*) with ROSA-DTA mice (RRID:IMSR_JAX:009669) as in (*5*). The lines were genotyped using the following primers: Cre Fw: ATT TGC CTG CAT TAC CGG TCG; Cre Rv: CCC CAG AAA TGC CAG ATT ACG TAT ATC; R26 DTA Fw: CAT CAA GGA AAC CCT GGA CTA CTG; R26 WT Fw: AAA GTC GCT CTG AGT TGT TAT; R26 Rv: GGA GCG GGA GAA ATG GAT ATG.

### Embryonic cortical 2D primary cultures

Somatosensory cortices from E16 mouse embryos were microdissected in ice-cold HBSS 1x (Gibco, cat#14025092), mechanically dissociated with TrypLE™ Express Enzyme 1x (Gibco, cat #12604013) after 5 minutes incubation at 37°C and collected by centrifugation (500 x g for 5 minutes, 4C). Dissociated cells were then filtered through a 70um cell strainer (VWR, 352350) and plated in 24-well plates containing Neurobasal medium pre-supplemented with Penicillin-Streptomycin (ThermoFisher, cat#15070063), 2% B27 supplement and 0.5mM Glutamine onto glass coverslips pre-coated overnight at 37°C with 0.1mg/ml Poly-D-Lysine (Advanced Biomatrix, cat#5174-5MG) and Laminin 0.01mg/ml (Thermo Fisher, cat#23017015). Cultures were kept in incubators at 37°C with 5% CO2 and partial replacement of the culture media was done every 2-3 days.

### Embryonic cortical organotypic cultures

Somatosensory cortices from E16 mouse embryos were microdissected in ice-cold HBSS 1x (Gibco, cat#14025092) and acute slices were obtained using Leica Vibratome VT1220S (cutting parameters: 0.1 mm/s speed, 1mm amplitude, 300 um thickness). Microdissected S1 cortices were subsequently cultured with the same medium of 2D cultures using organotypic cell culture inserts (Merck Millipore, PICM03050). Cultures were kept in incubators at 37°C with 5% CO2 and full media change was done once every 2-3 days.

### *In utero* electroporation

Timed pregnant CD1 mice were pre-emptively injected with 300ul of Buprenorphine 0.1 mg/kg (Temgesic, Benckiser, Switzerland) in 0.9% NaCl, and subsequently anaesthetized with isoflurane (5% induction, 3% during the surgery). Embryos were injected unilaterally with 700nl of plasmid solution, diluted in endotoxin-free TE buffer and 0.002% Fast green FCF (Sigma), in the lateral ventricle. Embryos were electroporated by placing the head between circular tweezer-electrodes (5mm diameter, Sonidel Limited, UK), while 5 electric pulses (30 V for E12.5, 40 V for E14.0, 50 V for E15.5, 50 ms at 1 Hz) were delivered with a square-wave electroporator (Nepa Gene, Sonidel Limited, UK).

### Plasmids

The following plasmids were used for *in utero* electroporations: pAAV-CAG-tdTomato (Addgene #83029); pCAG-GFP (Addgene #16664); pAAV-CAG-ChR2(E123T;T159C)-2A-tDimer (Addgene #85399); pCAG-ChR2-Venus (Addgene #15753).

### Single-nuclei RNA sequencing (snRNA-seq)

#### SnRNA-seq of mouse somatosensory cortices

Somatosensory cortices were collected for E17 embryos, P1, P7, P14 CD1 mice. For Reeler model, at P23, three brains were collected from three independent litters of Reln^rl+/+^ mutants and corresponding Reln^rl+/-^ control littermate. VB- somatosensory cortices were collected from three Slc6a4-Cre; LSL-DTA mutant mice, and from three LSL-DTA^+/-^ control mice at P14 from the same litter. For E17 time point, somatosensory cortices from 5 embryos were microdissected in ice-cold HBSS 1x (Gibco, cat#14025092). For postnatal timepoints, three to four-hundred-micrometer-thick coronal slices were prepared and primary and secondary somatosensory cortices were microdissected. The tissue was preserved at -80°C until use for the single nuclei suspension, which was obtained through douncer homogenization (Sigma D8938) using an adaptation of the protocol for the Nuclei EZ buffer kit (Sigma, NUC101).

For dissociations, after being thawed on ice, the tissue was resuspended in 2mL of ice-cold extraction buffer (Nuclei EZ prep buffer) and dounce-homogenized on ice until disintegration (15 to 25 pestle A and B). After incubation for 5 minutes in 4ml EZ buffer total, the extracted nuclei were collected by centrifugation (500 x g at 4°C, 5 minutes). The extraction procedure was repeated once, before performing a washing step, in which nuclei were resuspended in washing buffer composed of 1% BSA in PBS, with 50U/ml of SuperASE-In (Thermo Fisher, cat# AM2696) and 50U/ml of RNasin (Promega, cat# N2611), after centrifugation. Finally, the nuclei were resuspended in 1mL of washing buffer and filtered through a 30μm strainer into a new low-binding 1.5mL tube and processed for fluorescence-activated nuclei sorting (FANS) on Beckman Coulter MoFlo Astrios, after incubation with Hoechst (Invitrogen H3570, 1:500) for 5-10 minutes on ice. 20k nuclei were sorted and 42ul of nuclei suspension was used to load the Chromium Next GEM Single Cell 3’ Kit v2 (10x Genomics), according to the manufacturer’s instructions.

### SnRNA-seq from primary cultures

Single-nuclei suspensions were obtained as indicated in the manual of the Nuclei EZ prep (cat#NUC-101) and nuclei isolation was performed through FANS as indicated for in vivo preparations. SnRNA-seq preparations were performed as indicated in manufacturers’ instructions.

### SnRNA-seq from organotypic cultures

Nuclei were harvested from cortical organotypic slices by douncers homogenization (pestle A 7 times and pestle B 6 times) by Nuclei EZ buffer extraction, similarly to tissue preparations. Nuclei were isolated through FANS and snRNA-seq preparations were performed according to manufacturers’ instructions.

### Analysis of snRNA-seq data

All single-nucleus transcriptomics data were mapped, and count matrix generated using 10x Genomics cellranger program (v6.1.1). The count matrices were then analyzed using the R statistical Software (v4.1.1), Seurat package (v5.0.1), ggplot (v3.3.6), DoubletFinder (v2.0.3), clusterProfiler (v4.10.1), bmrm (v4.4), vegan (v2.6-6.1), bayestestR (v0.13.2).

#### Cells filtering and quality control

The data of CD1 mice *in vivo, in vitro* and organotypic slices cultures, transcriptomes were considered as single nuclei when expressing more than 500 genes, showing less than 30000 UMI counts, which UMI/gene ratio was superior at 1.2 and percent of mitochondrial RNA lower than 2%. The Reeler mutant and control P23 transcriptomes were filtered for expressing 1000 genes and more, UMI counts between 2500 and 15000 and a percent of mitochondrial RNA lower than 1%. The VB mutant and control transcriptomes were filtered for expressing more than 500 genes, expressing less than 20000 UMIS and less than 2% of mitochondrial RNA. In addition to quality controls, cells annotated as microglia, endothelial and vascular-related identities were excluded from further analysis for all datasets.

#### Cell population hierarchy

Public single cell transcriptomics dataset of the entire adult mouse neocortex and hippocampus(*11*) was used to build the hierarchy of cell populations. Cells harvested from the somatosensory primary cortex were isolated and endothelial and vascular cell types (Endo, Micro-PVM, SMC-Peri and VLMC) along with Cajal-Retzius (CR) cells were removed from further analysis. Cells were then analyzed following the Seurat standard procedure. The data was clustered using the *FindClusters()* function from Seurat with different values of the resolution parameter corresponding to 0.0005 for cell classes, 0.001 for subclasses and 0.08 for types (Fig. S1A).

#### Selection of population marker genes

The three reference datasets (CD1 P14, DTA^flox/flox^ P14 and Reeler^+/-^ P23) were merged and integrated using CCA integration from Seurat v5 pipeline. Following integration, cells were annotated to cell types, subclasses and classes using a Deep Neural Network trained to classify cells based on the Yao et al. 2021 dataset as described previously(*28*). The code and details of the DNN (Deep Neuronal Network) training are available at: https://github.com/baumannnatalia/Celltype_Classifier_DNN. The marker genes were identified using the *FindMarkers()* function from Seurat, with minimal log2FC value of 0.6 and minimal difference of percentage of expressing cells of 0.1.

#### Identity UMAP generation

All count matrices were scaled using the *SCTransform()* function from the Seurat package. The average gene expression of the different population markers was measured as the mean of scaled expression of all genes of a given population. The averaged population gene expression was then used for UMAP visualization. All reference dataset cells used for marker identification were used as reference. For each cell, the 10 nearest reference neighbors were identified using Euclidean distance on the UMAP 2D space. The class, subclass and type identities of the nearest neighbors were evaluated to assign a cell to a certain population. If 9 or 10 neighbors belonged to the same population, the cell was annotated to this population. If the neighbors’ identities were mixed (less than 9 neighbors within the same population), the cell was considered undefined. Each hierarchical level was annotated independently, meaning a cell could be assigned to a population for higher hierarchical level but not for the lower ones (Fig. 1D).

#### Differentially expressed genes, gene ontologies and KEGG pathway enrichment

The differentially expressed genes shown in Fig. S3 were measured using the *FindMarkers()* function from the Seurat package. The genes were filtered to have a log fold change higher than 0.6, an adjusted p-value lower than 0.05 and a difference in percentage of expressing cells higher than 0.2. The enrichment analysis of gene ontologies within differentially expressed genes was done using the *enrichGO* function for biological processes from the clusterProfiler package. The 10 most enriched gene ontologies were grouped into larger function classes annotated by hand and shown on Fig. S3. The KEGG pathway enrichment was performed with *enrichKEGG* from the clusterProfiler package. The top 10 enriched pathways are shown in the dotplot and the top 5 in the network of Fig 5J.

#### Pseudo-age prediction

Pseudo-age was built using ordinal regression models from bmrm package as previously described(*29*). Briefly, a regularized ordinal regression was used order cells harvested at different time points from the *in vivo* dataset (E17, P1, P7, P14). During modeling, a weight is assigned to each gene to optimize the correct ordering of cells. To avoid overfitting, a cross-validation was performed to assess the performance of the modeling. To retrieve the optimal number of genes for the pseudo-age model, different numbers of genes were tested. The efficiency of the models was assessed by measuring the overlap of distributions of the cross-validation values of cells harvested at different time points, using the *Overlap* function from the bayestestR package. A high overlap coefficient means poor separation of cell from different time point, we selected the minimal number of genes for which the overlap coefficient was minimal, here 180 genes (Fig. S3 H-I, Fig. S4D). The pseudo-age value of *in vivo* cells used for model training corresponds to their cross-validation value, while for the 2D *in vitro* and organotypic cells, the prediction values are shown.

#### Shannon diversity index

To assess the diversity of cell population from each class and in each environmental conditions, all data from *in vivo* E17 to P14, *in vitro* DIV1 to DIV17 and organotypic DIV10 and DIV17 were merged (without integration). A clustering analysis was performed using the *FindClusters* function from the Seurat package with resolution parameter set to 1. The Shannon diversity index was measured from the number of cells within each cluster using the *diversity* function from the vegan package with the index option set to “shannon”.

### Immunofluorescence and imaging

#### Immunofluorescence staining of 2D primary culture

Coverslips were fixed with 4% PFA in PBS for 15 minutes at room temperature and washed three times in PBS. Blocking and permeabilization were performed by incubating the cultures, at room temperature, with 1% BSA, 0.5% Triton-X in PBS for 45 minutes. Afterwards, incubation with the primary antibodies was done overnight at 4°C in antibody solution (0.5% BSA, 0.1% Triton-X in PBS). Following three washes in PBS, the coverslips were incubated at room temperature for 1 hour with secondary antibody mix and Hoechst, and finally mounted on glass slides, after being washed three times in PBS. Images were acquired on Nikon A1r or LSM800 confocal microscopes (Carl Zeiss).

#### Immunofluorescence staining of organotypic culture

Slices were fixed with 4% PFA in PBS for 30 minutes at room temperature and washed three times in PBS. Blocking and permeabilization were performed by incubating the cultures, at room temperature, with 1% BSA, 0.5% Triton-X in PBS for 45 minutes. Afterwards, incubation with the primary antibodies was done overnight at 4°C in antibody solution (0.5% BSA, 0.1% Triton-X in PBS). Following three washes in PBS, the coverslips were incubated at room temperature for 1 hour with secondary antibody mix and Hoechst, and finally mounted on glass slides, after being washed three times in PBS. Images were acquired on Nikon A1r or LSM800 confocal microscopes (Carl Zeiss).

The following antibodies were used: chicken anti-MAP2, 1:5000 Abcam ab5392; goat anti-PDGFRa 1:500 Bio-Techne AF1062; rabbit anti-GFAP 1:1000 Abcam ab7260; rabbit anti-DCX 1:500 Cell Signaling 4604; mouse anti-GAD67 1:500 Merck Millipore MAB5406; rabbit anti-β3 tubulin 1:1000 BioLegend PRB-435P.

#### Electrophysiology & optogenetics

For *ex vivo* patch-clamp connectivity recordings, three-hundred-micrometer-thick coronal slices were prepared from P14 to P21 CD1 mice electroporated at E14 or E15.5 with pCAG-Venus-ChR2 plasmid. Slices were kept for 30 min in artificial cerebrospinal fluid (ACSF) at 33°C (125 mM NaCl, 2.5 mM KCl, 1 mM MgCl2, 2.5 mM CaCl2, 1.25 mM NaH2PO4, 26 mM NaHCO3 and 11 mM glucose, saturated with 95% O2 and 5% CO2) before recording. The slices were then transferred in the recording chamber, submerged and continuously perfused with ACSF.

For *in vitro* patch-clamp connectivity recordings, recordings were performed on cocultured neurons in utero-electroporated neurons at E14 with pCAG-Venus-ChR2 plasmid and electroporated at E15.5 with pCAG-tdTomato plasmid or in utero-electroporated at E15.5 with pCAG-Venus-ChR2 plasmid and electroporated at E14 with pCAG-tdTomato plasmid. The internal solution used for the experiments contained 140 mM potassium methansulfonate, 2 mM MgCl2, 4 mM NaCl 0.2 mM EGTA, 10 mM HEPES, 3 mM Na2ATP, 0.33 mM GTP and 5 mM creatine phosphate (pH 7.2, 295 mOsm). EPSC were evoked in presence of 1µM tetrodotoxin (Tocris, CAS Number 18660-81-6) and 1mM 4-aminopyridin (Hellobio, CAS Number 504-24-5) to avoid polysynaptic recording and 10µM bicuculine (Tocris, CAS Number 485-49-4) by a 10 msec blue-light pulse at 0.1 Hz delivered through the 60X objective. For resting membrane-potential and excitability recordings, immediately after the whole-cell configuration the neuron was placed in current clamp mode and 4 steps of 50pA, 100pA, 200pA and 400pA were applied for 500ms. Membrane potential was monitored every 10 s and averaged for 5 consecutive acquisitions, within the first 2 min after the whole-cell configuration establishment. Access resistance was monitored by a hyperpolarizing step of 14 mV at each sweep, every 10s. Liquid junction potential was not corrected. Recordings were amplified (Multiclamp 700, Axon Instruments), filtered at 5 kHz and digitalized at 20 kHz (National Instrument Board PCI-MIO-16E4, IGOR WaveMetrics), and stored on a personal computer for further analyses (IGOR PRO WaveMetrics).

## Acknowledgments

We thank Julien Guy from the Staiger lab for the Reeler mice, the Genomics Platform, Bioimaging, HPC and FACS Facility of the University of Geneva, A. Benoit for technical assistance, L. Frangeul for logistic support, all members of the Jabaudon laboratory, as well as the members of the Tole laboratory for constructive exchanges during the project.

## Authors contributions

S.F. and D.J. conceived the project, N.B., I.M, S.R.P, E.K., G.B., S.F. performed experiments, N.B. performed bioinformatic analyses, N.B, I.M., S.F and D.J. wrote the manuscript with support from all authors.

