## Supplementary Figures for "Cell-extrinsic controls over neocortical neuron fate and diversity"

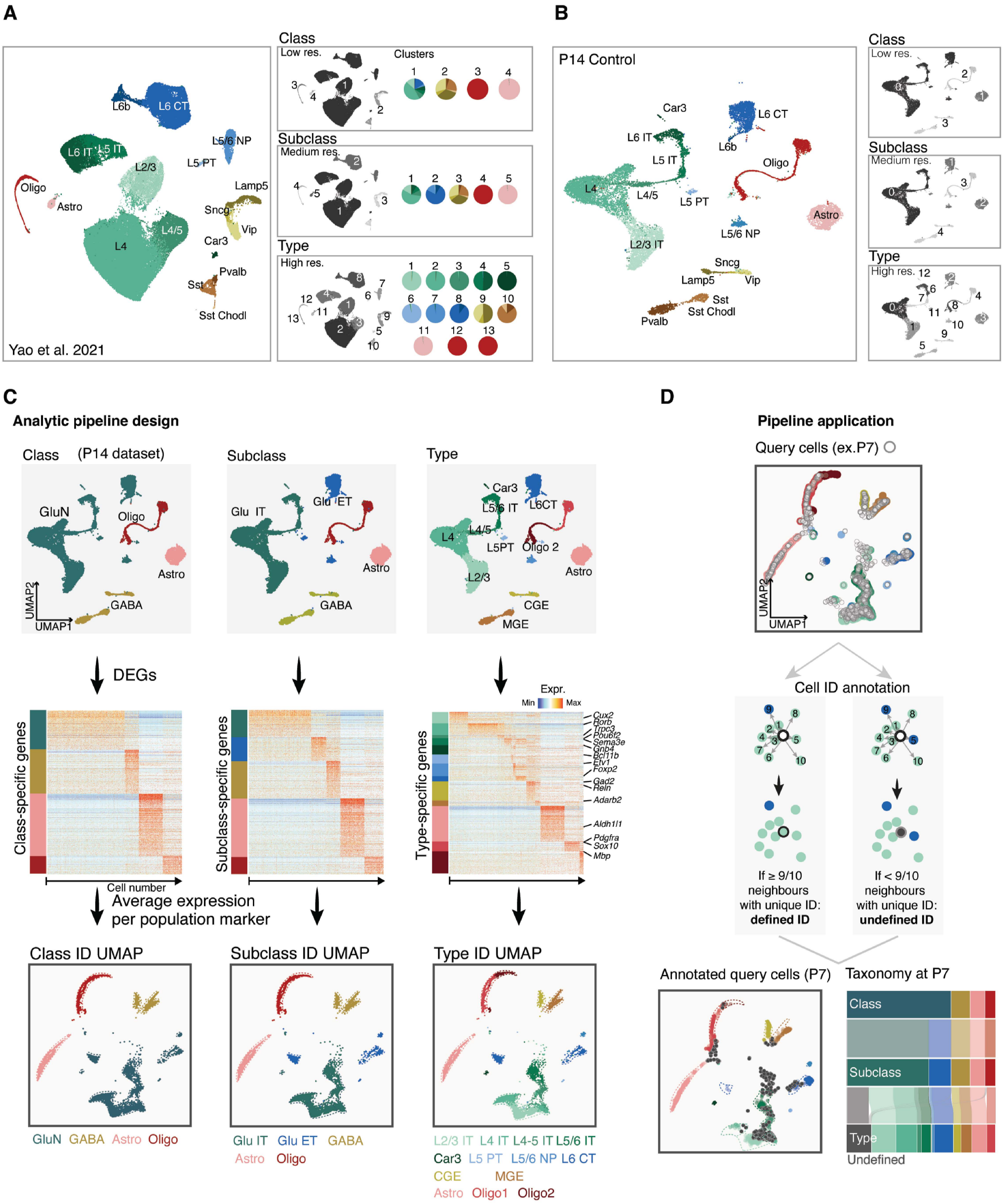


**Figure S1: Related to Figure 1, defining cell identity hierarchy.**
**A,** UMAP representation of the Yao et al, 2021 reference dataset used for the hierarchical tree construction. On the right, the three clustering resolutions used for the classification at in classes, subclasses and types are shown, with the pie charts showing the percentages for each population in the respective clusters. **B,** the same UMAP representations are shown for the reference P14 control used for the identity-based annotation. **C,** representation of the analytical pipeline used to generate the reference UMAP for each hierarchical level. On the top, the initial UMAPs, followed by the heatmaps showing markers expression for each population (DEGs), and at the bottom the resulting “identity UMAPs” at each hierarchical level. **D,** Schematic description of the analytical pipeline used to annotate cell identities for a query dataset (the example is for P7 in vivo dataset) and the resulting outcomes with Identity UMAP

(bottom left) and the hierarchical representation of the population identity annotations with the Sankey plot (bottom right). *Abbreviations*: Glu, glutamatergic neurons; IT, intratelencephalic neurons; ET, extratelencephalic neurons; Astro, astrocytes; Oligo, oligodendrocytes; L6 CT, layer 6 corticothalamic neurons; PT, pyramidal tract neurons; L2/3, layer 2/3; NP, near-projecting neurons; CGE, caudal ganglionic eminence-derived interneurons; MGE, medial ganglionic eminence-derived interneurons; ID, identity; DEGs, differentially expressed genes.

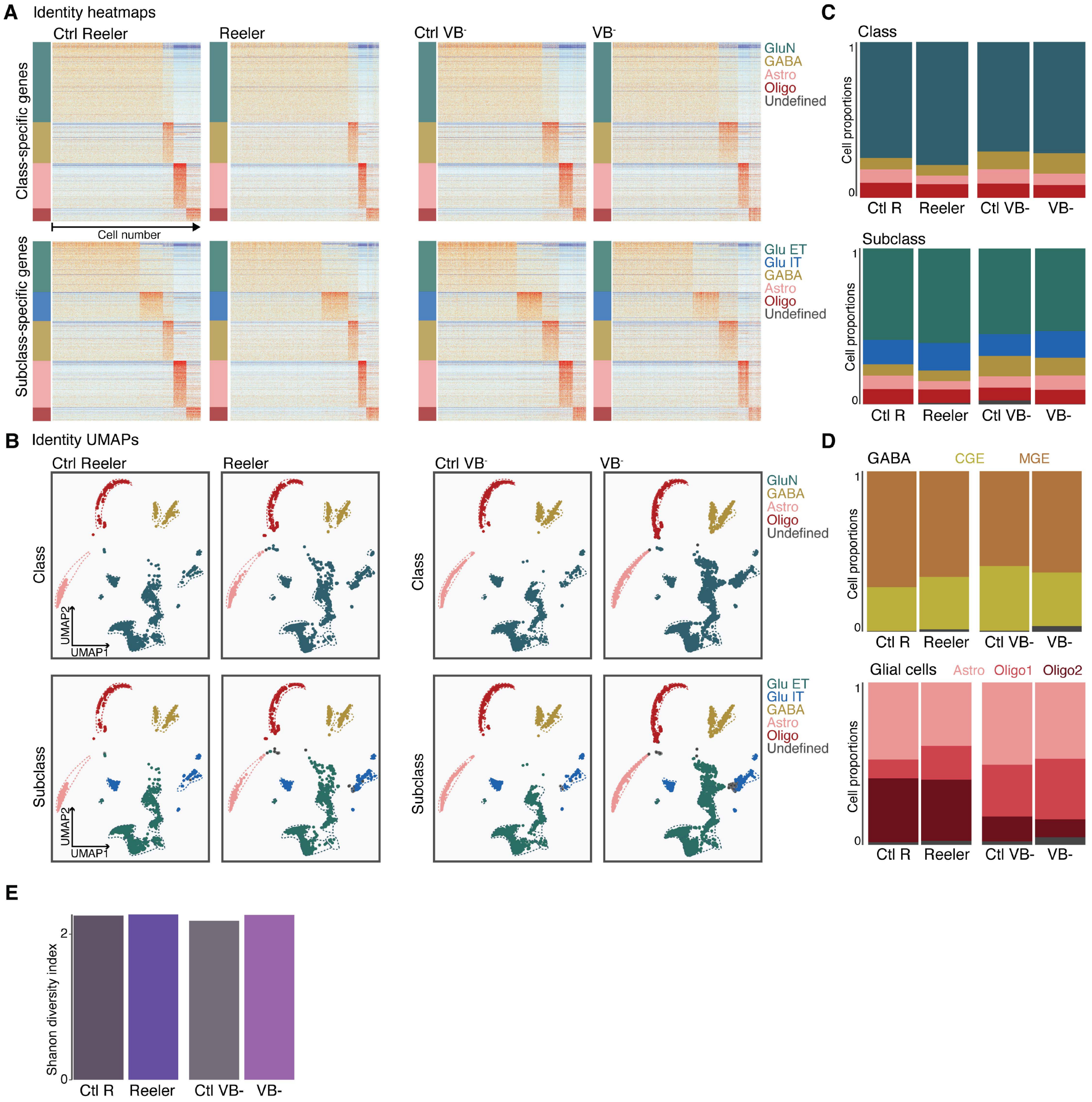


**Figure S2: Related to Figure 2, cell type-specific environmental sensitivity in Reeler and VB- mice.**

**A,** Identity heatmaps for cell classes and subclasses across the four conditions. **B,** Identity UMAPs for cell classes and subclasses across the four conditions. **C,** Proportions of cell classes (top) and subclasses (bottom) for the four analyzed conditions. **D,** Proportions of cell types within the GABAergic and glial cell classes for each of the four analyzed conditions. **E,** Shannon index indicating the cell type diversity for all four conditions. *Abbreviations*: Glu, glutamatergic neurons; IT, intratelencephalic neurons; ET, extratelencephalic neurons; Astro, astrocytes; Oligo, oligodendrocytes; CGE, caudal ganglionic eminence-derived interneurons; MGE, medial ganglionic eminence-derived interneurons.

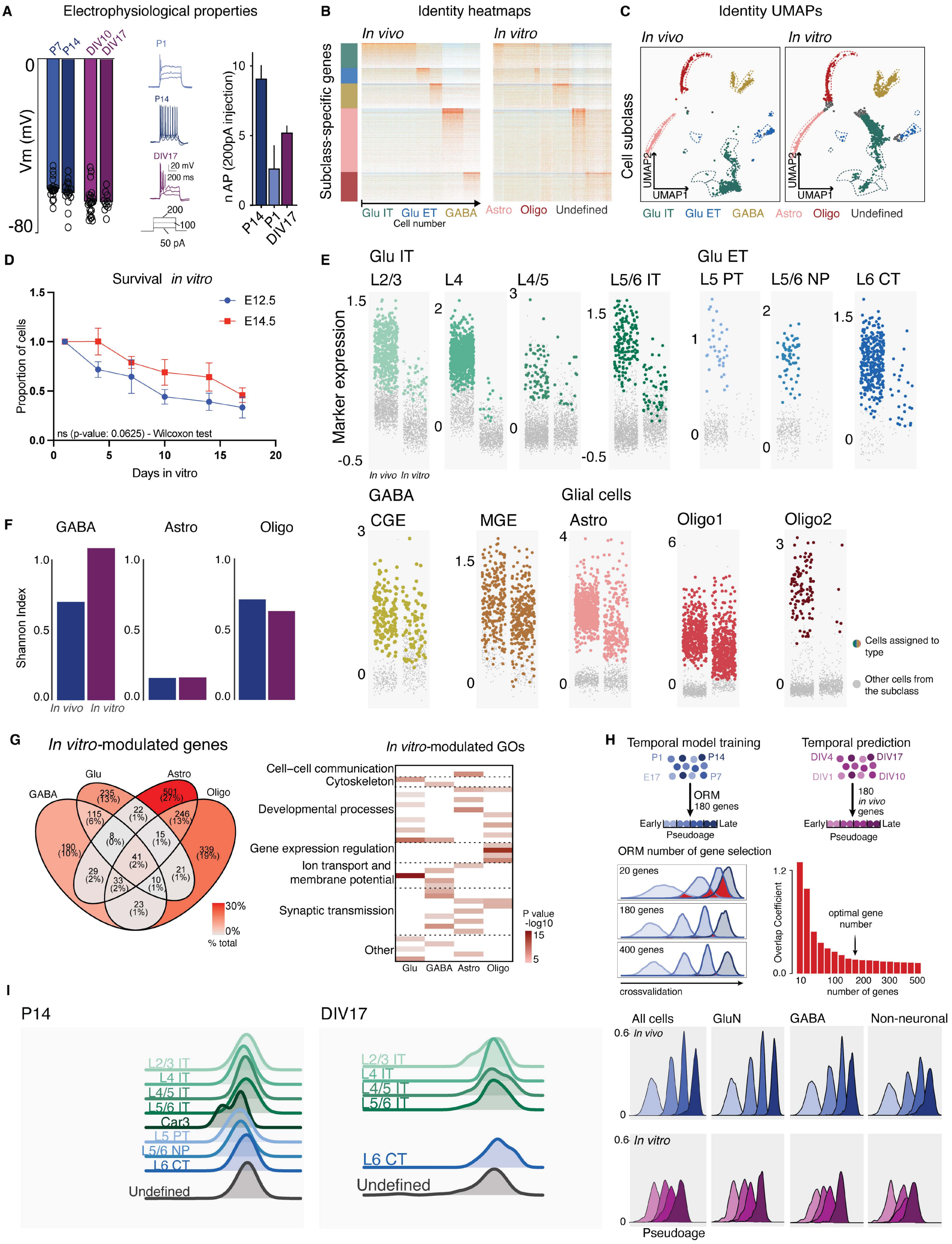


**Figure S3: Related to Figure 3, Cell population-specific molecular responses in 2D cultures.**
**A,** Electrophysiological measurements for neurons from P7 and P14 *in vivo* and DIV10 and DIV17 *in vitro*. Left, resting membrane potential values (Vm); middle, example traces for depolarizations; right, number of action potentials (APs) for P14, P1 and DIV17 showing excitability of the neurons. **B,** Identity heatmaps for cell subclasses for in vivo and in vitro conditions. **C,** Identity UMAPs for cell subclasses for in vivo and in vitro conditions. **D**, Survival

curves of E12.5 (blue) and E14.5 (red) electroporated neurons that were plated and followed longitudinally via confocal imaging. The y axis indicates the proportion of cells that survive in the culture over the days in vitro (x axis). The error bands represent standard error. **E,** Average markers expression for each indicated cell type (indicated above the graph) in the indicated cell populations for in vivo (left) and in vitro (right) conditions respectively. Each dot is one cell, and the colored cells were annotated while the grey dots represent cells that could not be annotated (undefined). **F,** Shannon index for GABA, astrocyte and oligodendrocyte cell classes for in vivo and in vitro conditions. **G,** Left, Venn diagram showing the overlap between differentially expressed genes between in vivo and in vitro calculated and P14 and DIV17 for each cell class. Right, heatmap for GO term enrichment for Biological Process for each cell class. **H,** Top, schematic of temporal model training using in vivo cells and its application for the pseudo-age prediction of in vitro cells. Bottom, pseudo-age curves for each of the timepoints in vivo and in vitro for all cells together (top left), or separately for the indicated cellular classes (top right and bottom). **I,** Temporal model prediction (left) for in vivo P14 condition and its prediction for the corresponding timepoint in vitro, DIV17, on the right for each cell type, including undefined cells at the bottom as separated group. The curves for some cell types are missing in vitro as their identities were undefined in this dataset. *Abbreviations*: Vm, resting membrane potential; nAP, number of action potentials; Glu, glutamatergic neurons; IT, intratelencephalic neurons; ET, extratelencephalic neurons; Astro, astrocytes; Oligo, oligodendrocytes; L6 CT, layer 6 corticothalamic neurons; PT, pyramidal tract neurons; L2/3, layer 2/3; NP, near-projecting neurons; CGE, caudal ganglionic eminence-derived interneurons; MGE, medial ganglionic eminence-derived interneurons; ORM, ordinal regression model; DIV, days in vitro; P postnatal days; ns, not significant; GO, gene ontology.

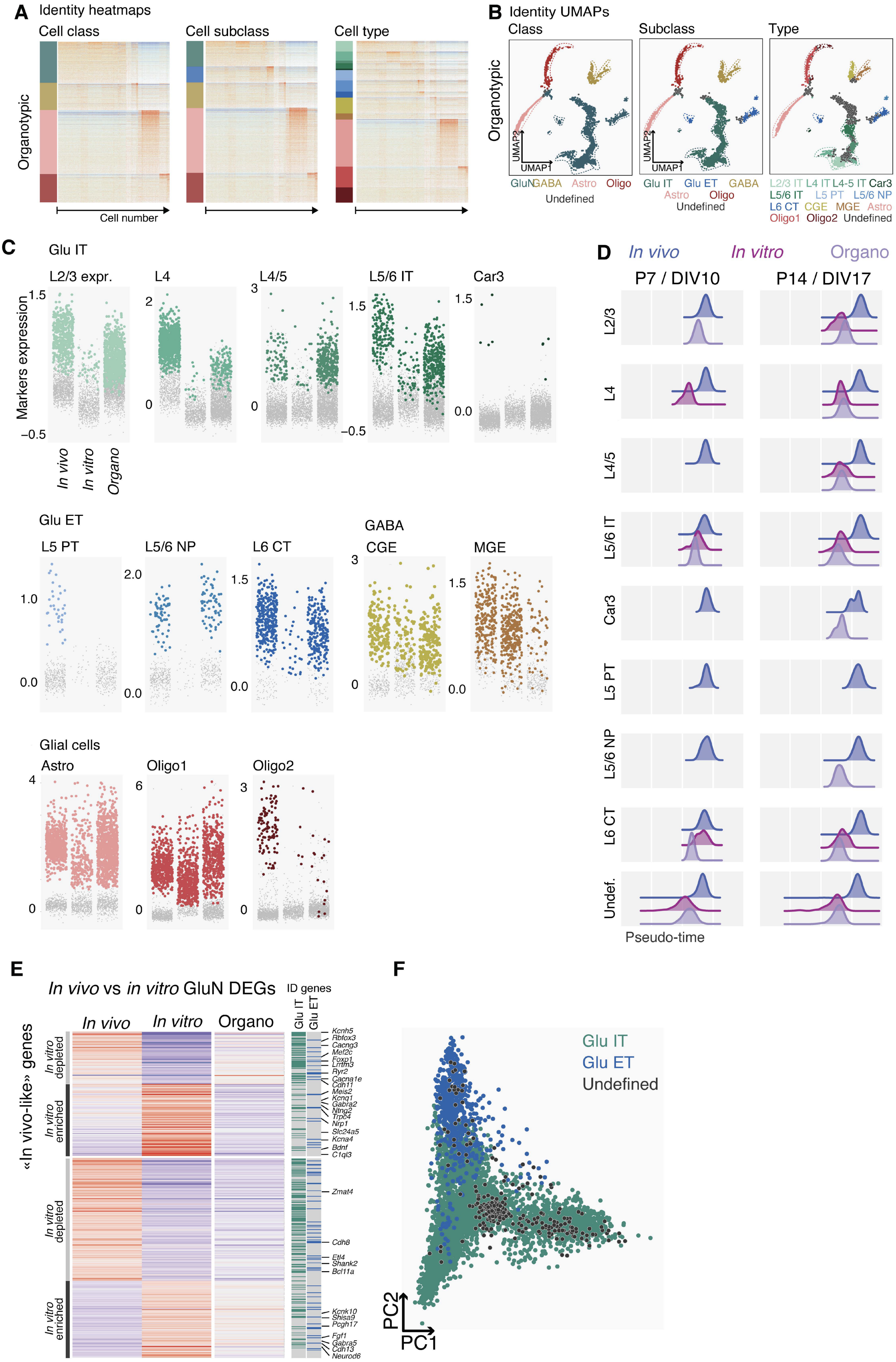
A Identity heatmaps
Cell class
Cell subclass
Cell type
Organotypic
Cell number
B Identity UMAPs
Class
Subclass
Type
Organotypic
UMAP1
UMAP2
GluN GABA Astro Oligo
Undefined
Glu IT Glu ET GABA
Astro Oligo
Undefined
L2/3 IT L4 IT L4-5 IT Car3
L5/6 IT L5 PT L5/6 NP
L6 CT CGE MGE Astro
Oligo1 Oligo2 Undefined
C
Glu IT
L2/3 expr.
L4
L4/5
L5/6 IT
Car3
Markers expression
In vivo
In vitro
Organo
Glu ET
L5 PT
L5/6 NP
L6 CT
GABA
CGE
MGE
Glial cells
Astro
Oligo1
Oligo2
D
In vivo
In vitro
Organo
P7 / DIV10
P14 / DIV17
L2/3
L4
L4/5
L5/6 IT
Car3
L5 PT
L5/6 NP
L6 CT
Undef.
Pseudo-time
E
In vivo vs in vitro GluN DEGs
ID genes
Glu IT
Glu ET
In vivo
In vitro
Organo
«In vivo-like» genes
In vitro depleted
In vitro enriched
Kcnh5
Rbfox3
Cacng3
Mef2c
Foxp1
Lrrtm3
Ryr2
Cacna1e
Cdh11
Meis2
Kcnq1
Gabra2
Ntng2
Trpc4
Nrp1
Slc24a5
Kcna4
Bdnf
C1ql3
Zmat4
Cdh8
Etl4
Shank2
Bcl11a
Kcnk10
Shisa9
Pcgh17
Fgf1
Gabra5
Cdh13
Neurod6
F
Glu IT
Glu ET
Undefined
PC1
PC2

**Figure S4: Related to Figure 5, Increased cell type-specific glutamatergic neuron identity definition in organotypic slices.**

**A,** Identity heatmaps at all hierarchical levels including all populations for the organotypic culture dataset. **B,** Identity UMAPs for all hierarchical levels including all populations for the organotypic culture dataset. **C**, Average marker expression for each indicated cell type (indicated above the graph) in the indicated cellular population (in bold on top of the graphs) for *in vivo* (left), *in vitro* (middle) and organotypic (right) conditions. **D**, Temporal model-predicted pseudo-age of organotypic (violet) and in vitro 2D (pink)-harvested cells compared to in vivo (blue) for each of the indicated cell types at the indicated time points (left, P7 and DIV10; right, P14 and DIV17). **E**, Top, heatmap showing the average expression level of all differentially expressed genes (rows) in vivo compared to in vitro, and their expression in organotypic culture in glutamatergic neurons. The values are displayed for each condition at the latest timepoint (P14 for in vivo and DIV17 for in in vitro 2D and organotypic). **F**, PCA plot related to Fig. 5J; here the dots, corresponding to single glutamatergic neurons, are color-coded per cell type. *Abbreviations*: Glu, glutamatergic neurons; IT, intratelencephalic neurons; ET, extratelencephalic neurons; Astro, astrocytes; Oligo, oligodendrocytes; L6 CT, layer 6 corticothalamic neurons; PT, pyramidal tract neurons; L2/3, layer 2/3; NP, near-projecting neurons; CGE, caudal ganglionic eminence-derived interneurons; MGE, medial ganglionic eminence-derived interneurons; DIV, days in vitro; P postnatal days.
